# Mechanism of histone H2B monoubiquitination by Bre1

**DOI:** 10.1101/2023.03.27.534461

**Authors:** Fan Zhao, Chad W. Hicks, Cynthia Wolberger

## Abstract

Monoubiquitination of histone H2BK120/123 plays multiple roles in regulating transcription, DNA replication and the DNA damage response. The structure of a nucleosome in complex with the dimeric RING E3 ligase, Bre1, reveals that one RING domain binds to the nucleosome acidic patch, where it can position the Rad6 E2, while the other RING domain contacts the DNA. Comparisons with H2A-specific E3 ligases suggests a general mechanism of tuning histone specificity via the non-E2-binding RING domain.

## Main

Post-translational modification (PTM) of histones plays a central role in regulating eukaryotic transcription. Monoubiquitination of histone H2B-K123 in yeast, K120 in humans (H2B-Ub), is a hallmark of actively transcribed genes that also plays a role in DNA replication, DNA repair and RNA processing^1-3^. H2B-Ub stimulates methylation of histone H3-K4 and K79, and recruits FACT (Facilitates Chromatin Transcription) to promote efficient transcriptional elongation^4,5^. Bre1 is a dimeric ubiquitin E3 ligase that targets the E2 ubiquitin conjugating enzyme, Rad6, to monoubiquitinate H2B-K123 in yeast^6,7^. In humans, the closely related RNF20/RNF40 heterodimer targets RAD6A/B to ubiquitinate histone H2B-K120^8,9^. Mutations and deletions of RNF20/40 are found in a variety of cancers and are indicators of poor prognoses^10^. The mechanism underlying the specificity of H2B-K123/120 ubiquitination is unknown.

We report here the cryo-EM structure of a Bre1 E3 ligase dimer bound to a nucleosome that reveals the molecular basis for specific ubiquitination of histone H2B. The fragment used in the study, Bre1 591-700, includes the RING domain and a coiled-coil that mediates dimerization^11^, and directs specific ubiquitination of H2B-K123^12^. Three distinct states of the Bre1-nucleosome complex were resolved at overall resolutions of 3.47 Å for state 1, 3.25 Å for state 2, and 3.21 Å for state 3 (Extended Data Figs.1,2 and Table 1). In each state, one Bre1 monomer was well ordered, with a local resolution of 3-4 Å, while the other monomer was resolved at resolutions ranging from 4Å to 6Å (Extended Data Fig. 2), indicating somewhat higher mobility. The well-resolved Bre1 density map enabled us to successfully model Bre1 dimer onto nucleosome using its crystal structure^11^. Although the complex contains a polyprotein fusion containing two copies of Bre1 and a C-terminal Rad6 protein joined by flexible linkers, Rad6 is not visible in our cryo-EM maps, presumably due to its weak and transient interactions with the Bre1 RING domain^13^ and the nucleosome^14^. Importantly, the fusion enzyme has even higher H2B-ubiquitinating activity on nucleosome as compared to the separated Bre1 and Rad6 enzymes (Extended Data Fig. 3), indicating that the linkers do not interfere with enzyme function.

**Table 1.**
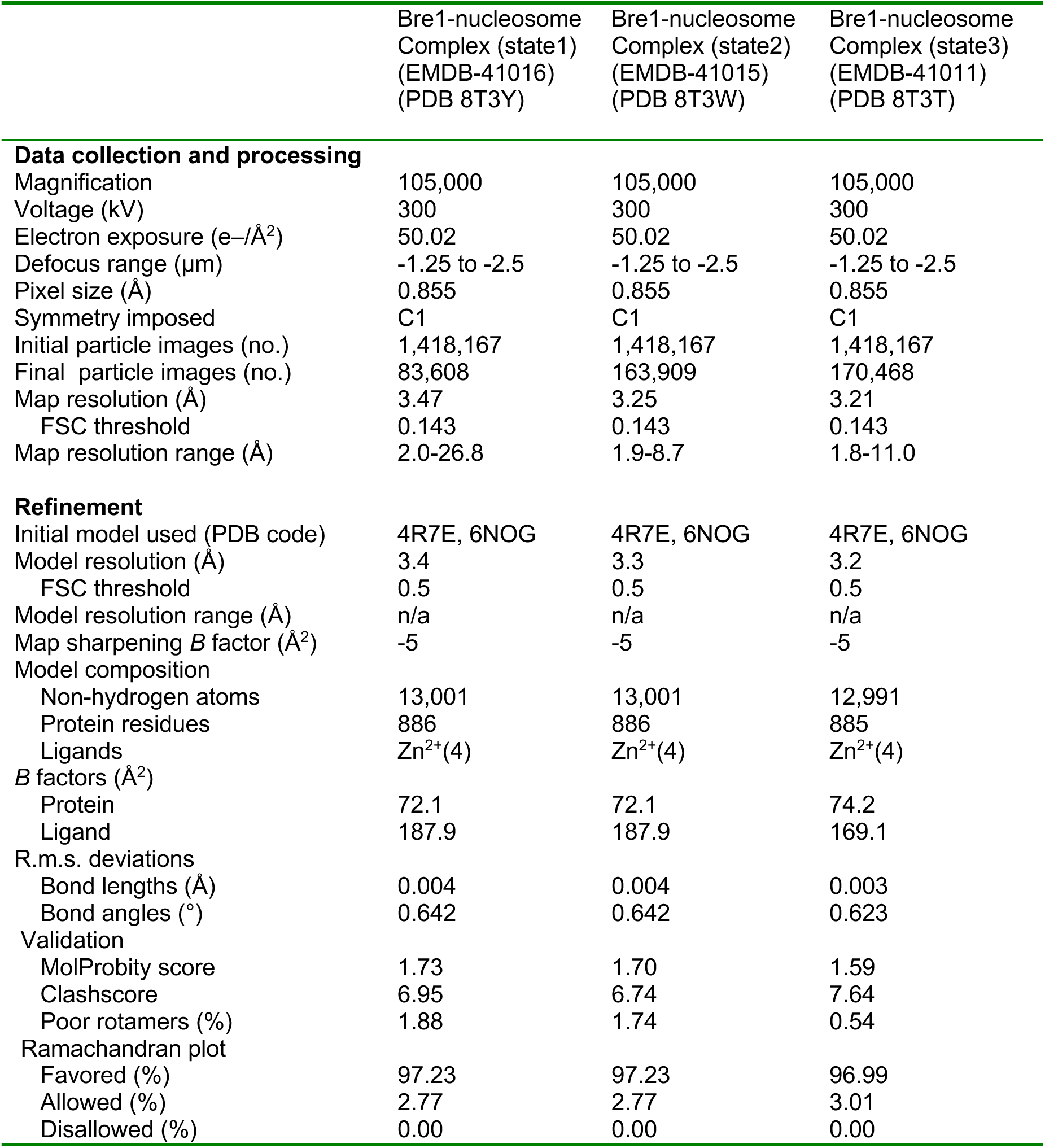
Cryo-EM data collection, refinement and validation statistics.

The Bre1 dimer straddles the periphery of the nucleosome, with one RING domain (Bre1-A) contacting the nucleosome acidic patch and the other (Bre1-B) interacting with the DNA at superhelical position (SHL) 6.5 (Fig.1a-b). Residues 632-647 form an α helix that mediates coiled coil interactions with the opposing monomer, followed by the catalytic RING domain, which contains a basic patch comprising residues R679, R681, K682, and K688 (Extended Data Fig. 4). The Bre1-A RING domain binds in a similar manner to the nucleosome acidic patch in all three states, with R679 forming salt bridges with H2A residues E61 and D90, and R681 forming salt bridges with H2A residues E61 and E64 (Fig.1b, c). This mode of interaction with the nucleosome acidic patch, termed an “arginine anchor,” has been observed in multiple structures^15^. The R679 contact represents a canonical arginine anchor while the R681 contact forms a type 2 variant arginine anchor^15^. The position of the Bre1-B RING relative to the DNA backbone varies more between the three states due to a repositioning of the Bre1 coiled coil. In state 1, the Bre1-B RING is positioned such that residues R679 and R681 are in a position to contact the electronegative DNA backbone (Fig.1d, Extended Data Fig.5a).

**Figure 1.**
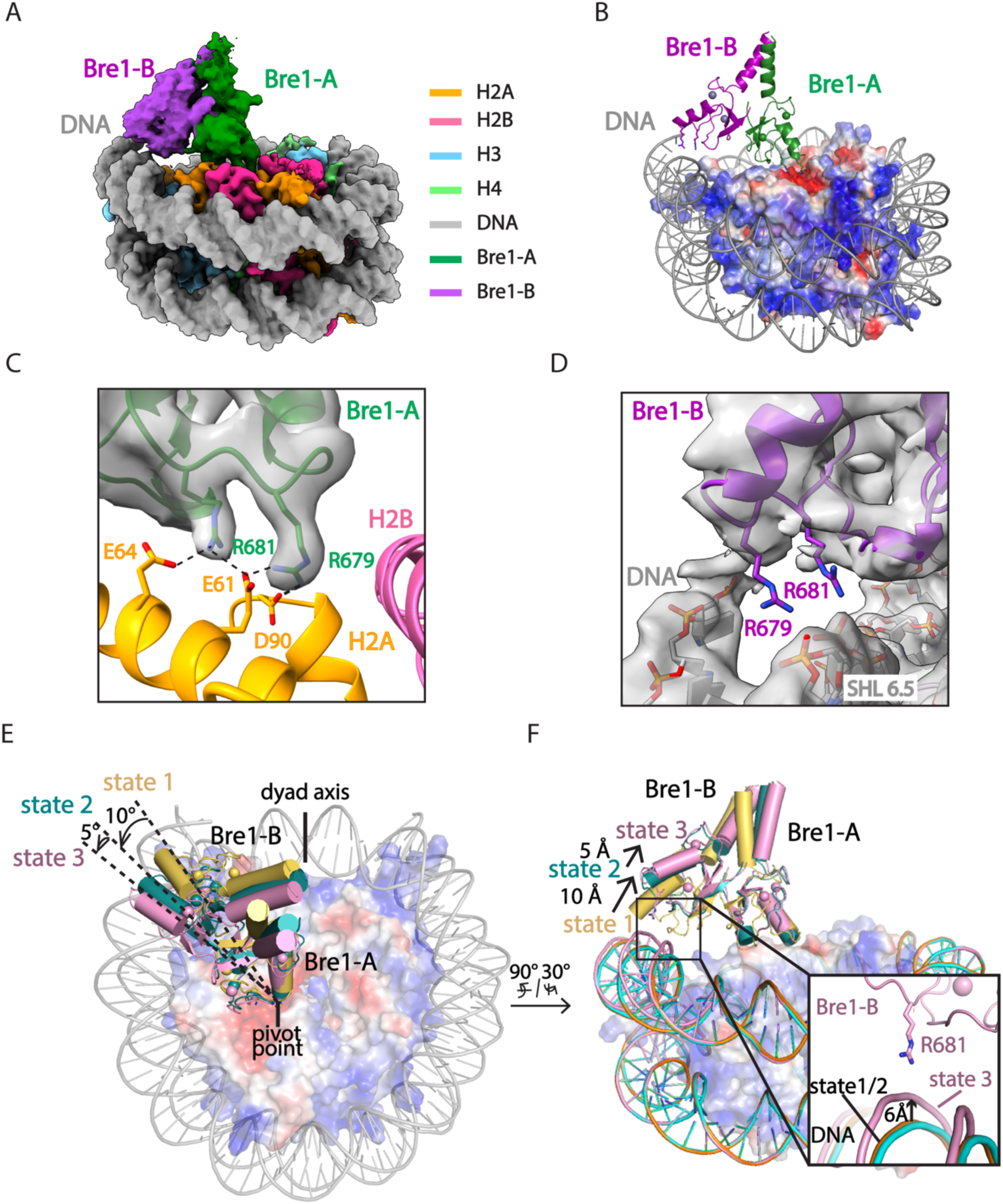
Cryo-EM structures of Bre1 dimer bound to nucleosome reveal multiple conformations. **a**, Cryo-EM density map of the Bre1 bound to nucleosome (state 1). **b**, View of the interface between Bre1 and nucleosome (state 1). Histone octamer is shown in an electrostatic potential surface representation (red=negative, blue=positive). **c**, Interface between Bre1-A (state 1) and the nucleosome acidic patch superimposed on EM map. **d**, Contact between Bre1-B and the DNA (state 1). The distances of Bre1-B R679 and R681 to DNA backbone are 3.4 Å and 4.3 Å. **e-f**, Change of Bre1 position in three states. The distance of Bre1-B R681 to DNA backbone in state 3 is 2.9 Å.

In state 2, the Bre1 dimerization coiled coil is tilted such that the Bre1-B RING rotates by about 10° about the face of the nucleosome and is about 10 Å farther from the DNA. In state 3, the Bre1 dimer is rotated by an additional 5° on the nucleosome relative to state 2, which shifts the Bre1-B RING by about 5 Å relative to state 2 (Fig.1e, f and Extended Data Fig.5). Importantly, there is a bulge in the DNA in state 3 that repositions the sugar-phosphate backbone 6 Å closer to the Bre1-B RING, placing Bre1-B residue R681 in a position to form electrostatic interactions with the DNA backbone (Fig.1f).

The key roles of Bre1 RING domain residues R679 and R681 in contacting the nucleosome acidic patch and the DNA are consistent with the deleterious effects of substitutions of these residues on Bre1 binding to nucleosomes and H2B ubiquitination activity (summarized in Table S1). Biochemical studies by Gallego et al.^14^ showed that Bre1 substitutions R679D and R681D abrogated nucleosome binding in pull-down assays and decreased H2B ubiquitination activity. Competition assays with the LANA (latency-associated nuclear antigen) peptide, which binds to the nucleosome acidic patch, showed that LANA peptide could compete with Bre1 for binding to the nucleosome acidic patch and reduce H2B monoubiquitination activity, further validates the importance of Bre1 contacts with the nucleosome acidic patch^14^. Importantly, an *in vivo* study conducted by Turco et al.^12^ showed that Bre1 basic patch mutations R675D/R679D and R681D/K682D completely abrogated H2B ubiquitination activity in yeast and resulted in severe growth defects comparable to a *bre1* deletion. Bre1 residues R679 and R681 that bind to nucleosome acidic patch are conserved in the RING domains of human homologues RNF20 and RNF40 (Extended Data Fig.6a) ^8^, suggesting a similar mechanism of H2B ubiquitination in humans.

The positioning of the E2, Rad6, by the Bre1-A RING domain relative to the H2B substrate lysine could be modeled based on the conserved manner of E2 enzyme binding to RING domains observed in crystal structures (Extended Data Fig. 7a)^16^ and on cryo-EM studies of Ubr1, a RING E3 ligase that also binds to Rad6 and polyubiquitinates N-degrons to target substrates for proteasomal degradation^17^. To model the position of Rad6 binding to the Bre1 RING, we aligned the Ubr1 RING domain to the Bre1-A RING domain for all three states of Bre1 bound to the nucleosome. Residues located at the predicted interface between the Bre1 RING and Rad6 are shown (Extended Data Fig. 7b-c). Of these, Bre1 substitutions L650A, R675D and K682D abolish H2B monoubiquitination in vitro^14^, while Bre1 mutants L650A and R675D lead to less ubiquitin discharge from Rad6∼Ub thioester, consistent with the predicted roles of these Bre1 residues in binding to Rad6 (Table S1)^14^.

A comparison of the Rad6 position in each state indicates that Bre1 alternates between active and inactive, poised states. In state 3, modeling positions the Rad6∼Ub thioester directly over histone H2B K120 (corresponding to K123 in the yeast nucleosome), with predicted distances of 2.2 Å between the Sγ of the Rad6 active site cysteine, C88, and Nζ of H2B K120, and 2.8 Å between the carbonyl carbon of Ub G76 and H2B K120 Nζ (Fig.2a, b). These distances would readily allow ubiquitin transfer from Rad6 to H2B K120. The corresponding distances in state 2 are 5.2 Å and 5.8 Å, respectively (Extended Fig. 8a-b). Interestingly, the modeled position of Rad6 in state 1 places the Rad6∼Ub thioester farther away from substrate lysine (8.8 Å and 10.7 Å) (Extended Data Fig. 8c, d). This suggests that each of the three Bre1-nucleosome complex states represents a different Rad6 activity status. Bre1 in state1 and 2 position Rad6 and Ub over the H2B K120 site, presumably in a “poised” state due to the distance from the substrate lysine, while state 3 is likely to facilitate an “active” state that promotes ubiquitin transfer to H2B K120. Interestingly, the histone H2A E3 ligase, BRCA1-BARD1, was also observed to adopt multiple orientations on the face of the nucleosome, with BARD1 forming distinct interactions with the C-terminal α-helix of histone H2B or with the H2B-H4 cleft ^18,19^.

**Figure 2.**
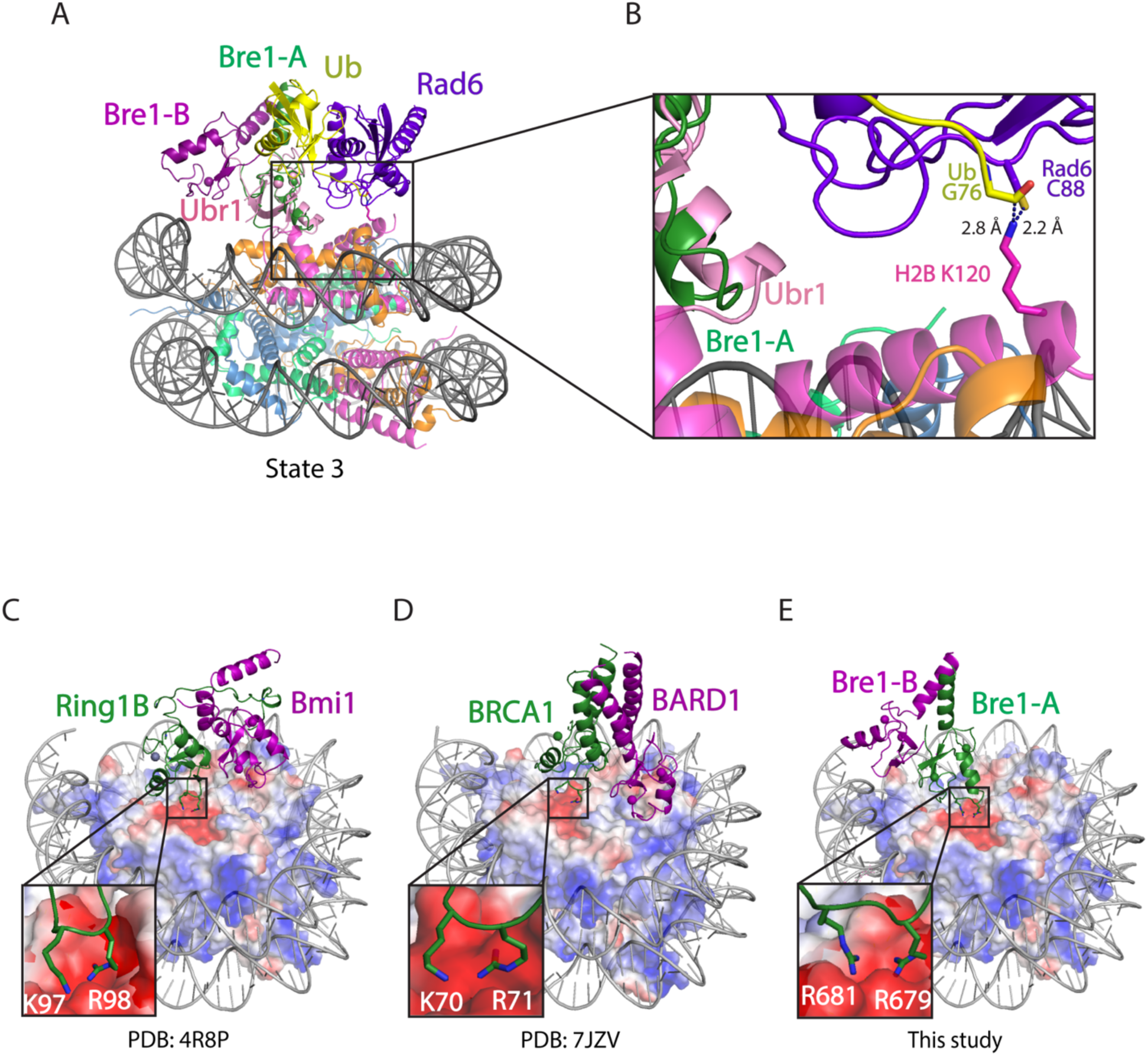
Bre1 orients Rad6 to ubiquitinate histone H2B K120. **a**, Modeling the position of Rad6 in Bre1-nucleosome state 3 by superposition of the Bre1 RING domain with Ubr1 RING-Rad6-ubiquitin (PDB: 7MEX) via the RING domain **b**, Close-up view of the modeled Rad6 active site C88, ubiquitin G76 and H2B K120. The H2B K120 rotamer with the shortest distance from Rad6 C88 and ubiquitin G76 is shown here. **c-e**, Structures of Ring1b-Bmi1 (PDB: 4R8P), BRCA1-BARD1 (PDB: 7JZV), and Bre1 dimer bound to nucleosome. Histone octamer is shown in an electrostatic potential surface representation.

Both Bre1 and Ubr1 contain N-terminal helical regions that contact Rad6 and enhance E2 discharge, termed RBD (Rad6 binding domain; residues 1-210) in Bre1 and U2BR (Ubc2-binding region) in Ubr1 ^12,17^. A recent crystal structure of Bre1-RBD bound to Rad6 revealed that the alpha-helical RBD forms a homodimer that binds to the Rad6 backside^20^. A superposition of the RBD-Rad6 complex with our modelled Rad6 bound to the Bre1-A RING shows that the RBD, which extends away from the nucleosome disk, can be readily accommodated without steric clash with the C-terminal Bre1 dimer or the nucleosome (Extended Data Fig.9). An additional ∼440 residues that are not present in the RBD or in the Bre1 fragment in the present study bridge the RBD C-terminus and the Bre1 coiled-coil.

Mutations in RNF20 and RNF40, have been found in a variety of cancers according to the cancer genomics database of cBioPortal (http://www.cbioportal.org). Of these, 27 map to residues in the RING domain or the α helical regions of RNF20 and RNF40 (Tables S2 and S3; Extended data Fig. 6a-b). RNF20 residue R955, corresponding to Bre1-R681 that binds the nucleosome acidic patch, is mutated to histidine in esophagogastric cancer (Extended data Fig.6a-b and Table S2). In addition, substitutions of RNF20 residues C924, R949 and Q954, which correspond to Bre1 residues L650, R675 and M680 that are predicted to interact with Rad6, are implicated in multiple cancers. Substitutions in these RNF20 residues that are predicted to disrupt the canonical E3 RING-E2 interface occur in non-small cell lung cancer (C924F, R949L, Q954*), cervical cancer (R949C) and colorectal cancer (R949H) (Extended data Fig.6a-b and Table S2).

A comparison of the Bre1 complex with that of E3 ligases that ubiquitinate other histone residues points to a pivot-like mechanism for tuning E3 ligase specificity. The positioning on the nucleosome of Bre1, which ubiquitinates H2B-K120/123, is markedly different from that of the heterodimeric E3 ligase, Ring1B/Bmi1^21^, which ubiquitinates H2A-K119, and BRCA1/BARD1^19^, which ubiquitinates histone H2A K125/127/129. Like Bre1, the RING domain that recruits the E2, Ring1B or BRCA1, binds to the nucleosome acidic patch with basic residues (Fig.2c-e). It is the orientation of the second RING domain in each complex that governs the positioning of the E2, and hence its specificity. Whereas Bre1 uses the second RING domain to interact with the DNA, Bmi1 and BARD1, respectively, bind to different locations in the globular histone core. Bmi1 caps the C-terminal end of H3 α1 helix via salt bridges formed by K62 and R64 and acidic residues in histones H3 and H4^21^, while BARD1 flexibly binds to the nucleosome H2B αC-helix^18^ or H2B/H4 cleft with its Trp91 inserted^19^. These second RING domains thus play the defining role in determining the E3 ligase specificity for its histone substrate.

## Acknowledgements

We thank Johns Hopkins colleagues Chuan Liu and Edward Twomey for advice on Cryo-EM data collection and processing and Xiangbin Zhang for valuable suggestions on constructing plasmids. Supported in part by the National Cancer Institute’s National Cryo-EM Facility at the Frederick National Laboratory for Cancer Research under contract 75N91019D00024. This work was supported by National Institute of General Medical Sciences grant GM130393 (C.W.).

## Contributions

C.W. and F.Z. conceived and designed the study. F.Z. and C.W.H. performed experiments, and processed data. F.Z. carried out modeling and map interpretation with assistance from C. W.. F.Z., C.W.H., and C.W. wrote the manuscript. C.W. supervised the study.

## SUPPLEMENTARY DATA

**Table S1.** Published effects of Bre1 mutants and LANA peptide competition assay.

| Mutation/expt | Description | Effect | Reference |
| --- | --- | --- | --- |
| Bre1-L650A | Bre1 RING domain contact with Rad6 | Defect in ubiquitin discharge from Rad6; Defect in H2B ubiquitination | Gallego et al, PNAS, 2016 |
| Bre1-R675D | Bre1 RING domain contact with Rad6 | Defect in ubiquitin discharge from Rad6; Defect in H2B ubiquitination | Gallego et al, PNAS, 2016 |
| Bre1-K682D | Bre1 RING domain contact with Rad6 | Defect in H2B ubiquitination | Gallego et al, PNAS, 2016 |
| Bre1-R679D | Arginine anchor in nucleosome acidic patch (Bre1-A); DNA binding (Bre1-B) | Defect in nucleosome binding and H2B ubiquitination | Gallego et al, PNAS, 2016 |
| Bre1-R681D | Arginine anchor in nucleosome acidic patch (Bre1-A); DNA binding (Bre1-B) | Defect in nucleosome binding and H2B ubiquitination | Gallego et al, PNAS, 2016 |
| Bre1-R675D/R679D | Contact with Rad6/ Arginine anchor in nucleosome acidic patch; DNA binding | Defect in nucleosome binding and H2B ubiquitination; Decreased H2B-Ub in yeast; Yeast growth defect | Turco et al JBC 2015 |
| Bre1-R681D/R682D | Arginine anchor in nucleosome acidic patch; DNA binding/Contact with Rad6 | Defect in nucleosome binding and H2B ubiquitination; Decreased H2B-Ub in yeast; Yeast growth defect | Turco et al JBC 2015 |
| Competition with LANA peptide | Binds acidic patch; standard test for acidic patch-binding proteins | Disrupts Bre1 binding and H2B ubiquitination | Gallego et al, PNAS, 2016 |

**Table S2.** RNF20 mutations located in RING domain and its preceded α helix.

| Study of Origin | Cancer Type | Protein Change | Mutation Type |
| --- | --- | --- | --- |
| Lung Squamous Cell Carcinoma (TCGA, PanCancer Atlas) | Non-Small Cell Lung Cancer | R949L | Missense_Mutation |
| Lung Squamous Cell Carcinoma (TCGA, PanCancer Atlas) | Non-Small Cell Lung Cancer | Q954* | Nonsense_Mutation |
| Uterine Corpus Endometrial Carcinoma (TCGA, PanCancer Atlas) | Endometrial Cancer | R971H | Missense_Mutation |
| Uterine Corpus Endometrial Carcinoma (TCGA, PanCancer Atlas) | Endometrial Cancer | D915Y | Missense_Mutation |
| Uterine Corpus Endometrial Carcinoma (TCGA, PanCancer Atlas) | Endometrial Cancer | K929N | Missense_Mutation |
| Uterine Corpus Endometrial Carcinoma (TCGA, PanCancer Atlas) | Endometrial Cancer | E907A | Missense_Mutation |
| Uterine Corpus Endometrial Carcinoma (TCGA, PanCancer Atlas) | Endometrial Cancer | L909M | Missense_Mutation |
| Lung Adenocarcinoma (TCGA, PanCancer Atlas) | Non-Small Cell Lung Cancer | C924F | Missense_Mutation |
| Skin Cutaneous Melanoma (TCGA, PanCancer Atlas) | Melanoma | H970Y | Missense_Mutation |
| Stomach Adenocarcinoma (TCGA, PanCancer Atlas) | Esophagogastric Cancer | P923L | Missense_Mutation |
| Stomach Adenocarcinoma (TCGA, PanCancer Atlas) | Esophagogastric Cancer | R955H | Missense_Mutation |
| Colorectal Adenocarcinoma (TCGA, PanCancer Atlas) | Colorectal Cancer | R949H | Missense_Mutation |
| Head and Neck Squamous Cell Carcinoma (TCGA, PanCancer Atlas) | Head and Neck Cancer | P923L | Missense_Mutation |
| Cervical Squamous Cell Carcinoma (TCGA, PanCancer Atlas) | Cervical Cancer | R949C | Missense_Mutation |

**Table S3.** RNF40 mutations located in RING domain and its preceded α helix.

| Study of Origin | Cancer Type | Protein Change | Mutation Type |
| --- | --- | --- | --- |
| Bladder Urothelial Carcinoma (TCGA, PanCancer Atlas) | Bladder Cancer | X944_splice | Splice_Site |
| Uterine Corpus Endometrial Carcinoma (TCGA, PanCancer Atlas) | Endometrial Cancer | R954C | Missense_Mutation |
| Uterine Corpus Endometrial Carcinoma (TCGA, PanCancer Atlas) | Endometrial Cancer | R954C | Missense_Mutation |
| Uterine Corpus Endometrial Carcinoma (TCGA, PanCancer Atlas) | Endometrial Cancer | R973W | Missense_Mutation |
| Uterine Corpus Endometrial Carcinoma (TCGA, PanCancer Atlas) | Endometrial Cancer | E937D | Missense_Mutation |
| Uterine Corpus Endometrial Carcinoma (TCGA, PanCancer Atlas) | Endometrial Cancer | A944T | Missense_Mutation |
| Uterine Corpus Endometrial Carcinoma (TCGA, PanCancer Atlas) | Endometrial Cancer | X944_splice | Splice_Region |
| Uterine Corpus Endometrial Carcinoma (TCGA, PanCancer Atlas) | Endometrial Cancer | A944V | Missense_Mutation |
| Uterine Corpus Endometrial Carcinoma (TCGA, PanCancer Atlas) | Endometrial Cancer | F967L | Missense_Mutation |
| Uterine Corpus Endometrial Carcinoma (TCGA, PanCancer Atlas) | Endometrial Cancer | R997H | Missense_Mutation |
| Stomach Adenocarcinoma (TCGA, PanCancer Atlas) | Esophagogastric Cancer | R973W | Missense_Mutation |
| Colorectal Adenocarcinoma (TCGA, PanCancer Atlas) | Colorectal Cancer | V966I | Missense_Mutation |
| Colorectal Adenocarcinoma (TCGA, PanCancer Atlas) | Colorectal Cancer | R973Q | Missense_Mutation |

**Extended Data Fig.1.**
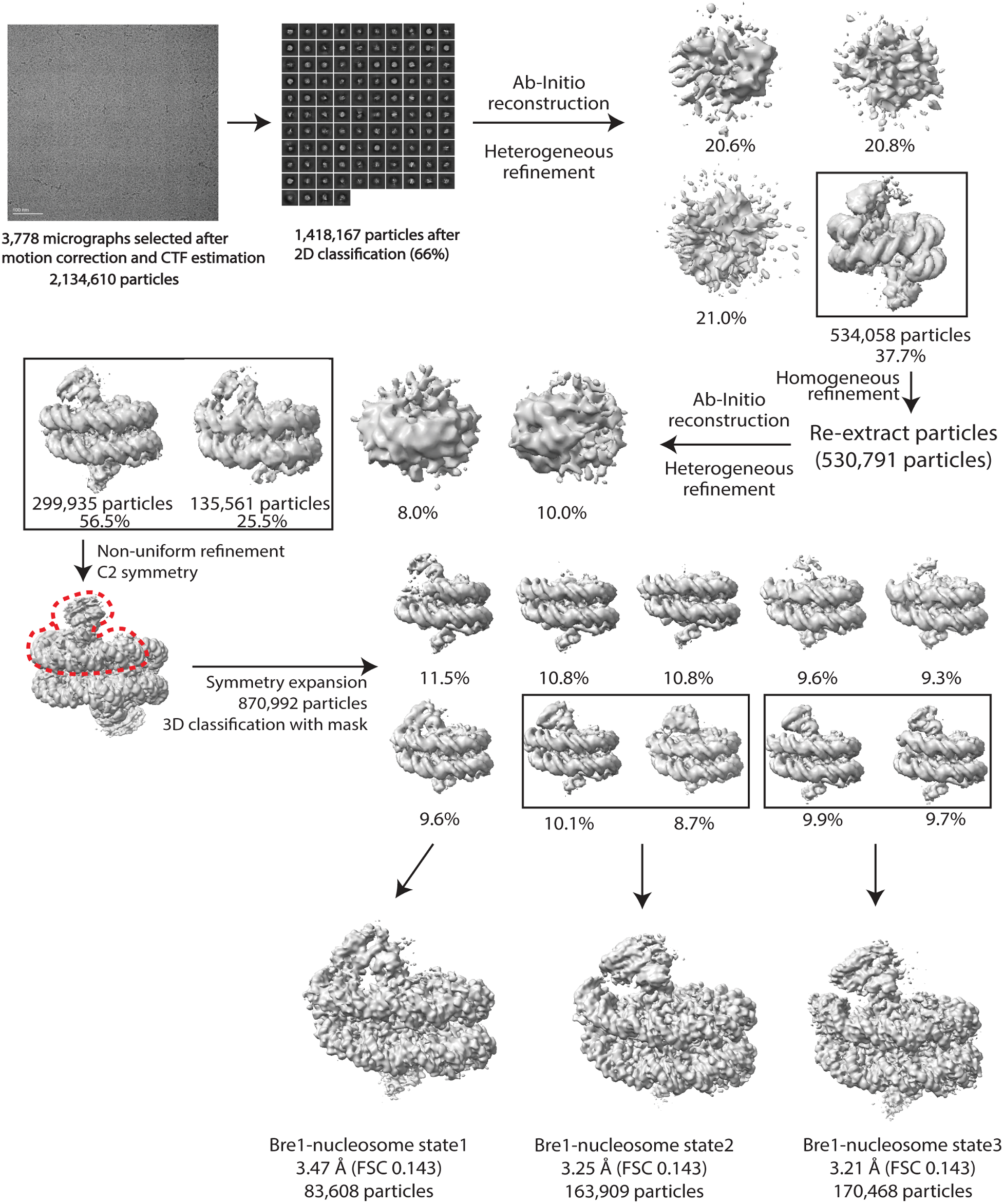
Cryo-EM processing workflow. The red dashed line indicates the mask generated for use in 3D classification. Three states of Bre1-nucleosome complex are identified and refined respectively.

**Extended Data Fig.2.**
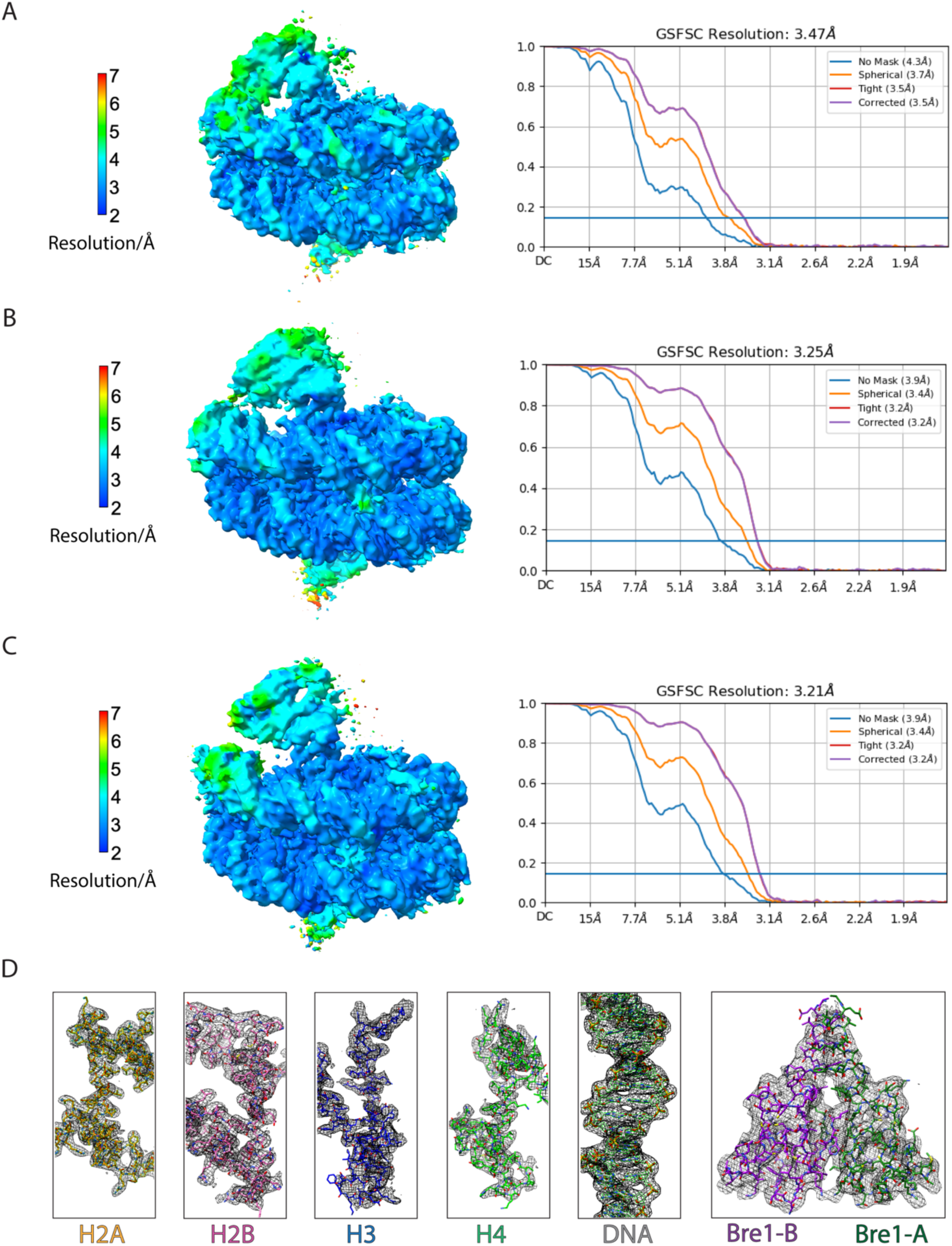
Cryo-EM validation and example density. **a-c**, Local resolution estimation and gold-standard Fourier shell correlation (GSFSC) curves at a FSC cutoff of 0.143 for each state of Bre1-nucleosome complex. **d**, Representative cryo-EM density maps for the histones, DNA, and Bre1.

**Extended Data Fig. 3.**
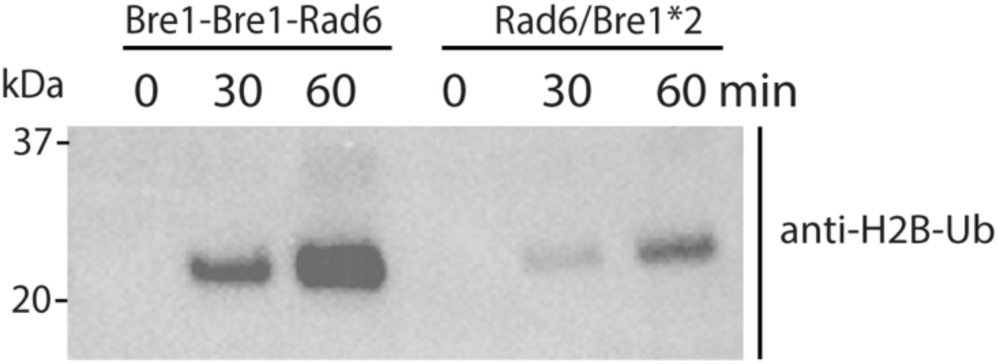
Comparison of enzymatic activity of fused and separated Bre1 and Rad6 enzyme. In vitro nucleosome H2B ubiquitination assay was performed with fused or separated Bre1 and Rad6 enzyme. Samples were quenched by SDS-loading buffer at indicated time points and subjected to immunoblotting. Data is representative of 2 independent experiments.

**Extended Data Fig.4.**
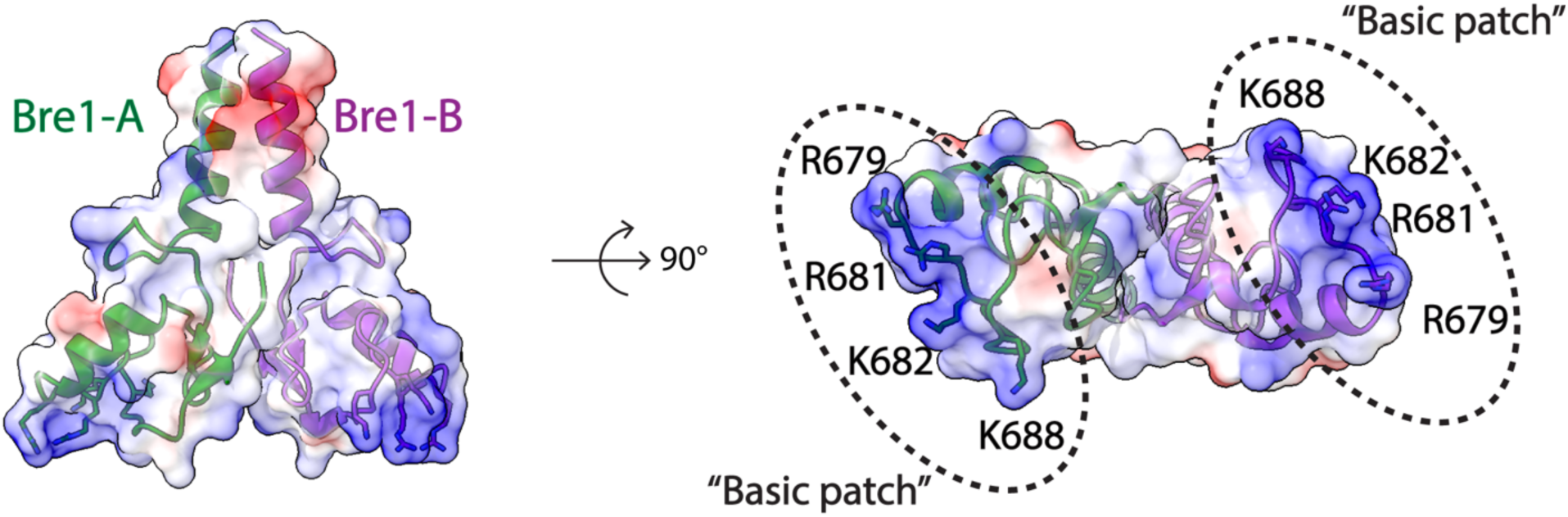
Electrostatic surface representation view of Bre1 dimer (PDB:4R7E). Bre1 is shown in a cartoon representation with basic residues highlighted as sticks.

**Extended Data Fig.5.**
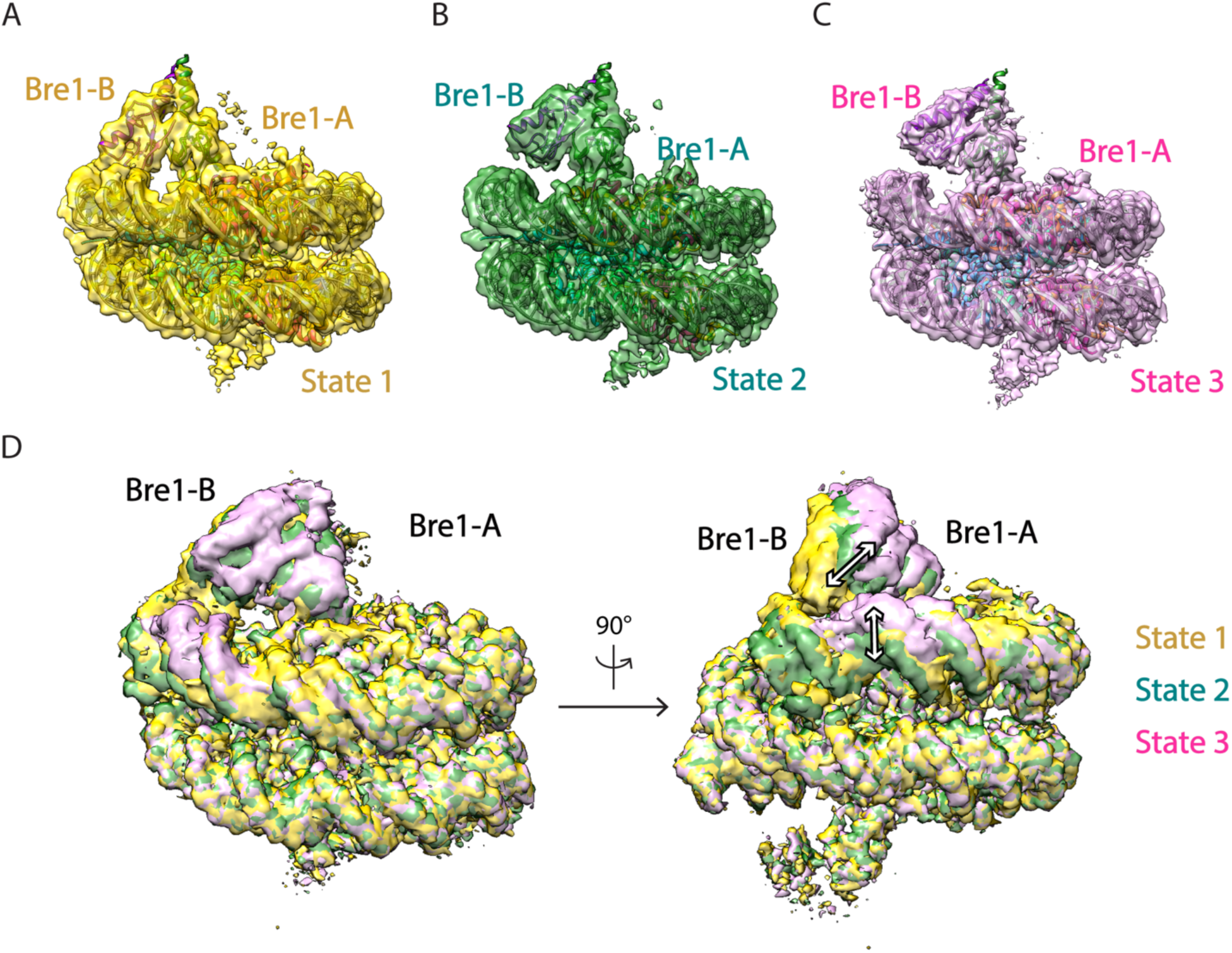
Multiple conformations of the Bre1-nucleosome complex. **a-c**, Cryo-EM maps of the Bre1-nucleosome complex with an atomic model fitted to states 1, 2 and 3. **d**, Superimpose of Bre1-nucleosome complex maps comparing states 1, 2 and 3. White arrows show the changes between each conformational state.

**Extended Data Fig.6.**
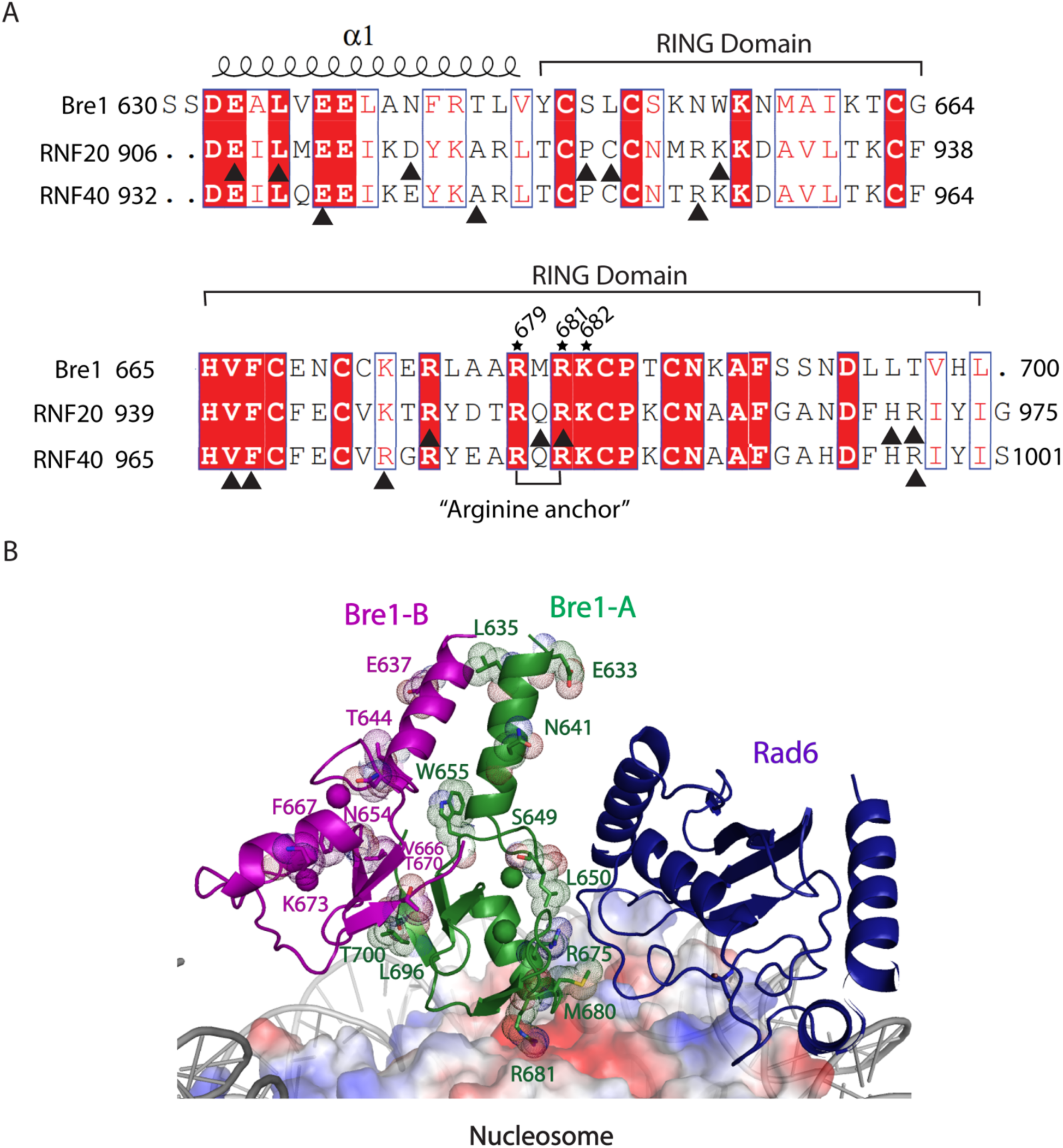
RNF20/40 mutation sites mapped to RING domain and preceded α-helix in cancer genomics. **a,** Sequence alignment of coiled coil and RING domains of Bre1, RNF20 and RNF40. Black triangle indicates the mutation sites of RNF20 and RNF40 in cancer genomics. **b**, Complex structure model of Bre1 RING-Rad6-nucleosome. Bre1 residues corresponding to RNF20/40 mutated sites were shown as sticks with dot sphere.

**Extended Data Fig. 7.**
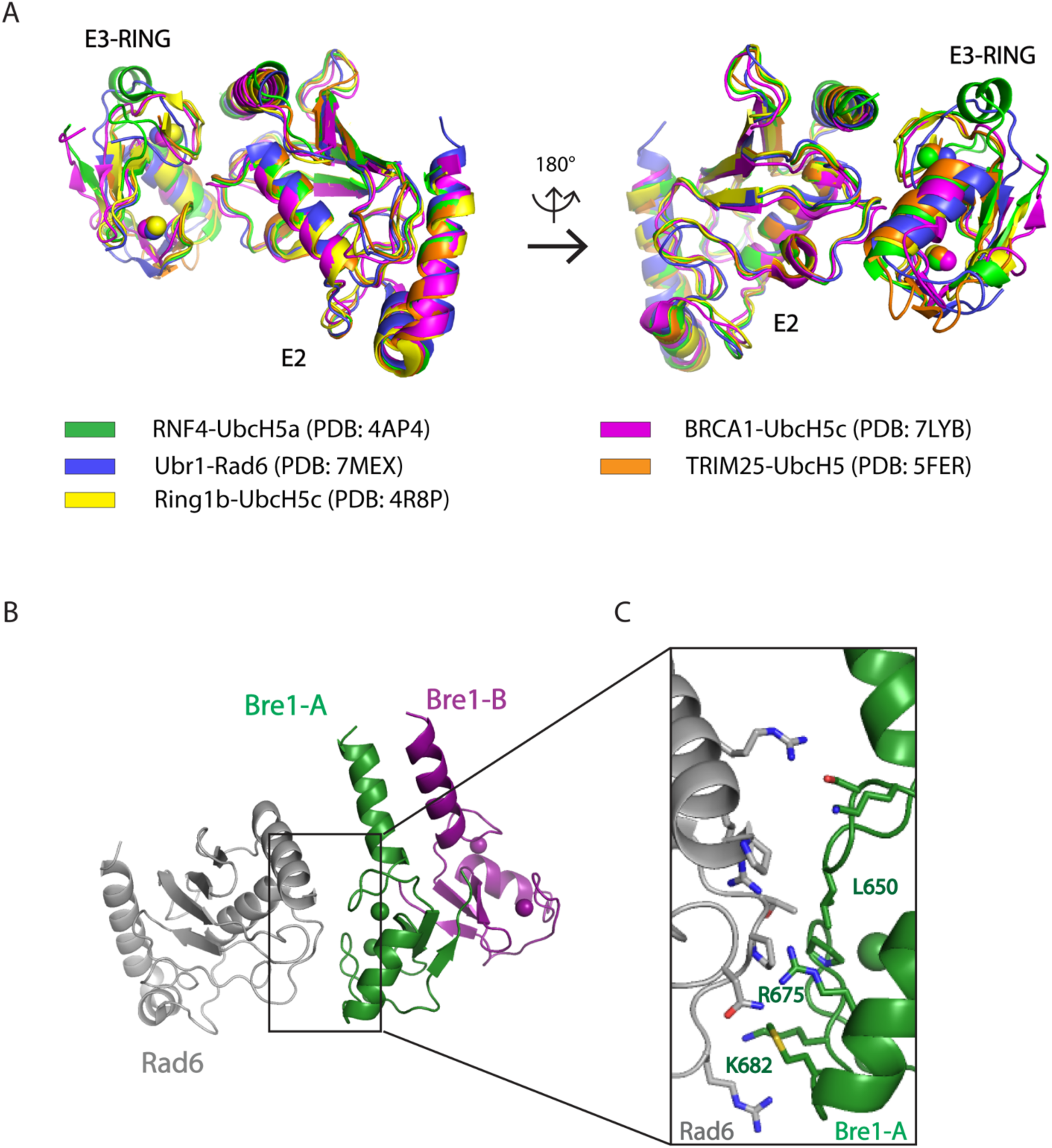
Canonical interface between E3 RING and E2. **a**, Superposition of multiple E2-E3 structures. E2-E3 complex structures of RNF4-UbcH5a (PDB: 4AP4), Ubr1-Rad6 (PDB: 7MEX), Ring1b-UbcH5c (PDB: 4R8P), BRCA1-UbcH5c (PDB: 7LYB) and TRIM25-UbcH5 (PDB: 5FER) are aligned together. **b**, The modelled structure of Bre1 RING-Rad6. **c**, Residues located in modelled Bre1A RING-Rad6 interface. The side chains of residues located in the interface are shown as sticks.

**Extended Data Fig. 8.**
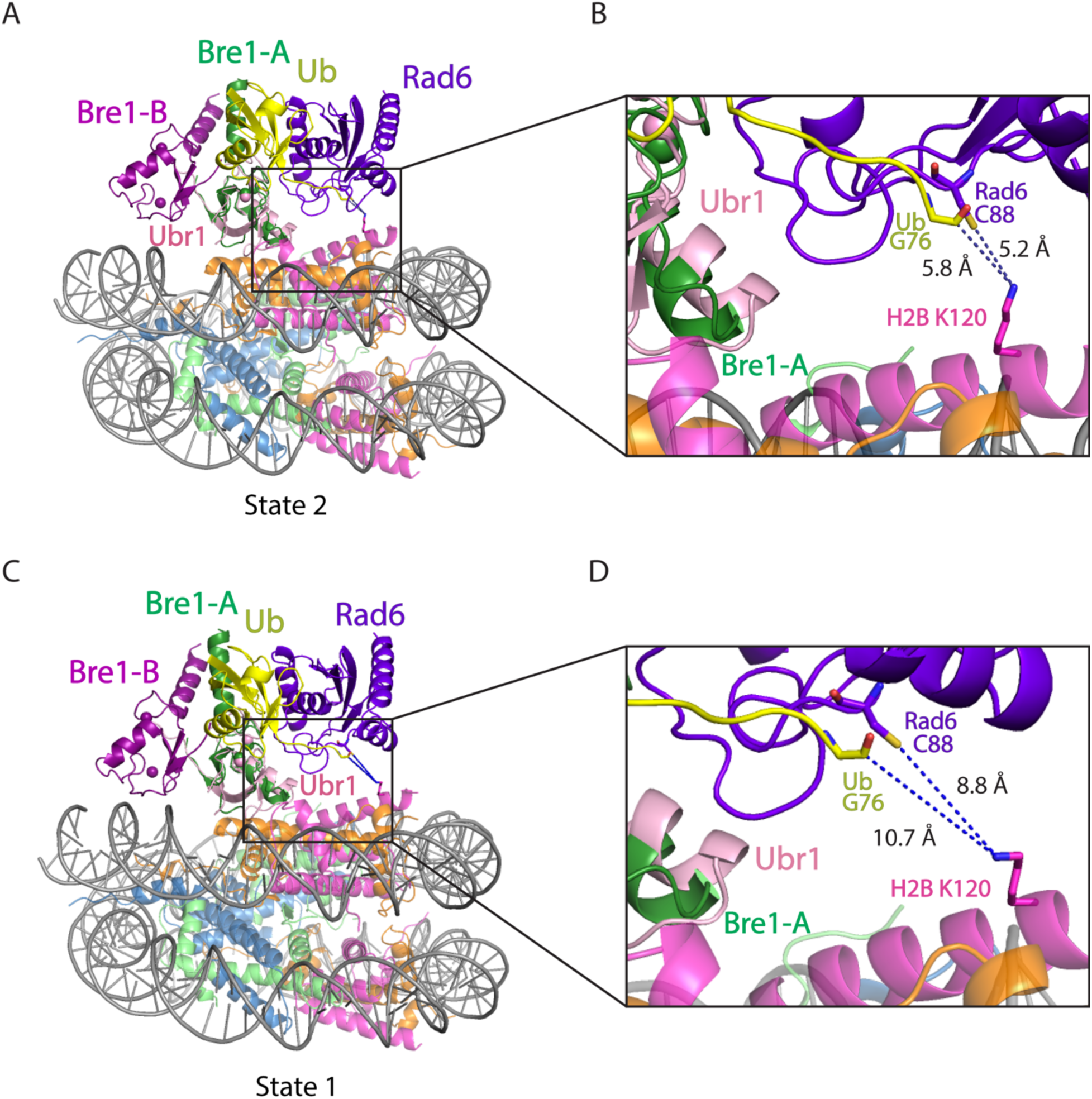
Superposition of Bre1-nucleosome complex (state 2 and 1) and Ubr1 RING-Rad6-ubiquitin (PDB: 7MEX). **a,c,** Ubr1 RING domain is aligned to Bre1-A RING domain in state 2 (a) and state 1 (c). **b,d**, Close-up view of Rad6 active site C88, ubiquitin G76, and H2B K120 in state 2 (b) and state 1 (d). The H2B K120 rotamer with the shortest distance from Rad6 C88 and ubiquitin G76 is shown here.

**Extended Data Fig. 9.**
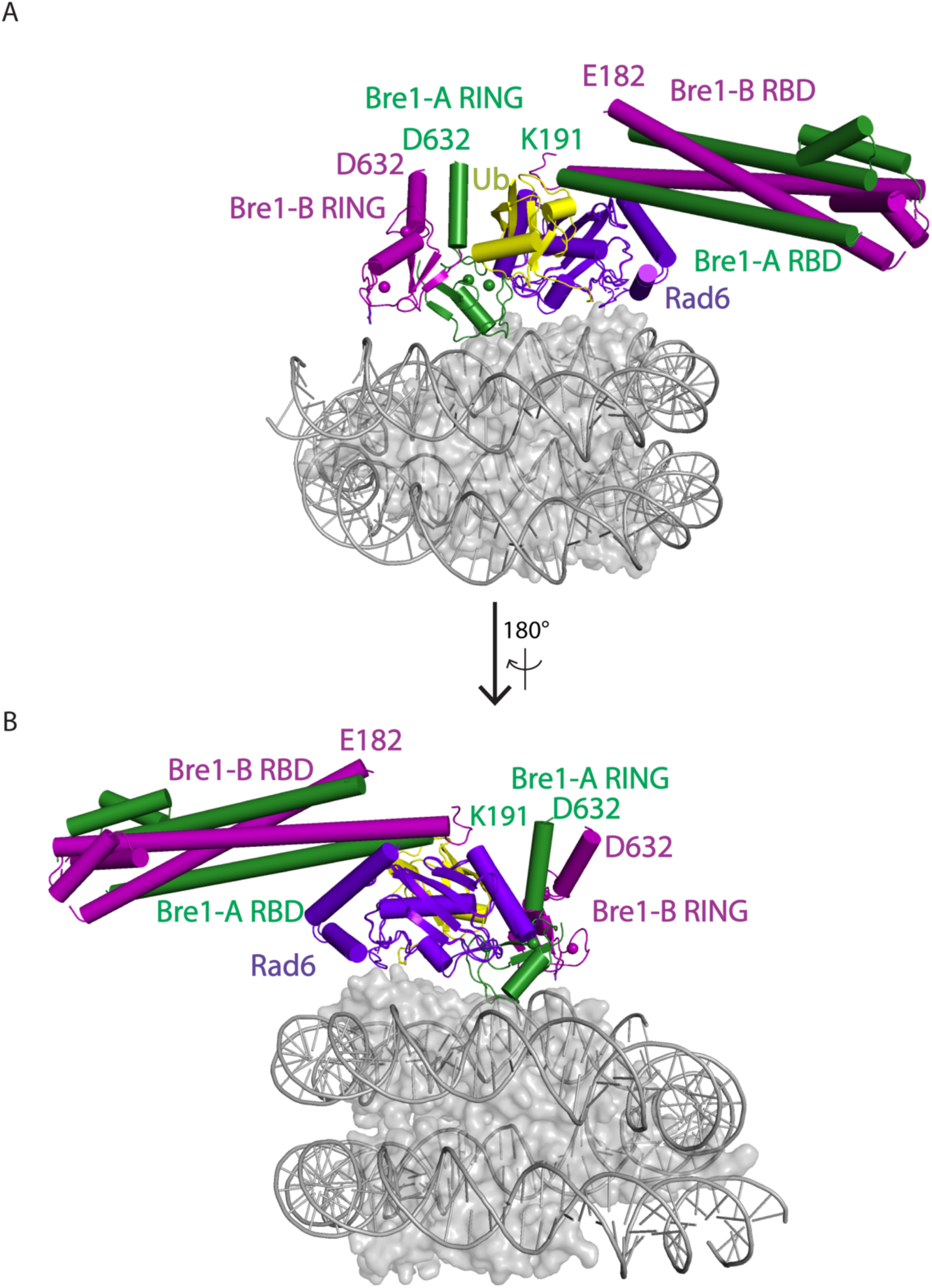
Structural alignment of Bre1 RBD-Rad6 complex and Bre1 RING-Rad6-Ub-nucleosome complex (model of state 3). **a, b,** Rad6 in Bre1 RBD-Rad6 complex is aligned to the Rad6 in Bre1 RING-Rad6-Ub-nucleosome complex (model of state 3).

## METHODS

### Protein expression and purification

A polymerase chain reaction (PCR) product of fusion protein Bre1(591-700) -(GGS)_4_- Bre1(591-700) -(GGS)_5_-Rad6(1-172) was cloned into vector pET32-a (Novagen) with N-terminal thioredoxin, a hexahistidine tag (Trx-His), and a Tobacco Etch Virus (TEV) protease site. This plasmid was used to transform BL21(DE3)Rosetta2 *E.coli* strain for protein expression. Briefly, *E. coli* cells were grown in Luria-Bertani (LB) media at 37°C while shaking until the culture reached an OD_600_ of 0.8. The growth temperature was then decreased to 18°C, 0.2 mM IPTG was added to induce protein expression, and the cells grown overnight.

Cells were harvested by centrifugation and resuspended in 20 ml lysis buffer (50 mM Tris pH 7.5, 300 mM NaCl, 50 μM ZnCl_2_, 5% glycerol, 5 mM 2-mercaptoethanol, 10 mM imidazole, 100 μM Phenylmethylsulfonyl fluoride (PMSF)) per liter of growth medium. Cells were lysed with a Microfluidizer (Microfluidics) and the cell lysate cleared by high-speed centrifugation. The supernatant was then incubated with 10 mL Ni-NTA beads (QIAGEN) for 90 min at 4°C. The beads were further washed with lysis buffer and the protein eluted with 50 ml lysis buffer supplemented with 250 mM imidazole. A 2 mg sample of TEV protease was added to the Ni-NTA column eluent for overnight cleavage. The resulting protein was further purified using a 5mL HiTrap Heparin column (Cytiva) developed with a salt gradient of 100 mM – 600 mM NaCl over 75ml followed by gel filtration on a Superdex 200 10/300 column (GE Healthcare) in buffer 20 mM HEPES, pH 7.5, 150 mM NaCl, 50 μM ZnCl_2_, 1 mM DTT. The purified protein was concentrated to 10 mg/mL, aliquoted, flash frozen, and stored at −80°C for future use.

### Nucleosome reconstitution

Unmodified *Xenopus laevis* histone proteins, H2A, H2B, H3 and H4 were purified as described previously^22^. The pST55−16 × 601 plasmid containing 16 repeats of the 147 base pair Widom 601 positioning sequence was amplified in the *E. coli* strain XL1-Blue. After plasmid extraction, the 601 DNA was excised with *Eco*RV, and recovered as described previously^23^. Nucleosomes were reconstituted from purified histones and DNA as previously described^23^.

### Cryo-EM sample preparation

A sample of 100 nM unmodified nucleosome was incubated with 10 µM Bre1-Bre1-Rad6 fusion protein in cross-linking buffer (20 mM HEPES pH 7.5, 50 mM NaCl, 1 mM DTT) at room temperature for 30 min. An equal volume of 0.15% glutaraldehyde was then added to the sample mixture. After incubation on ice for 1 hour, the cross-linking reaction was quenched by the addition of 100 mM Tris pH 7.5. The sample was dialyzed overnight into quenching buffer (50 mM Tris pH 8.0, 50 mM NaCl, 1 mM DTT) and applied to a Superdex 200 10/300 size exclusion column (GE Healthcare) that was pre-equilibrated with cross-linking buffer (20 mM HEPES pH 7.5, 50 mM NaCl, 1 mM DTT). The peak fractions were concentrated to a final sample concentration of 1 mg/mL.

Quantifoil R2/2 copper 200 mesh grids (Electron Microscopy Sciences) were glow-discharged for 45 seconds at 15 mA, after which, 3 µl of cross-linked sample was applied to the grids and blotted for 3.5 seconds with a blot force of 5, then immediately plunged frozen in liquid ethane using a Vitrobot Mark IV apparatus (Thermo Fisher Scientific) at 4°C, 100% humidity.

### Cryo-EM data processing and model building

A cryo-EM dataset was collected at the National Cryo-Electron Microscopy Facility (NCEF) of the National Cancer Institute on a Titan Krios at 300 kV utilizing a Gatan K3 direct electron detector equipped with an energy filter with a 20 eV slit width. A multi-shot imaging strategy (3 shots per hole) was used for data collection. Images were recorded in counting-mode at a nominal magnification of 105,000, a pixel size of 0.855 Å and a dose of 50.02 e^-^/Å^2^ with 40 frames per movie at a defocus range of 1.25-2.5 µM. A total of 4,561 movies were collected.

The dataset was processed using cryoSPARC^24^. After patch-based motion correction and contrast transfer function estimation, 3,778 micrographs were selected for particle picking. Initial particles were picked by Blob picker. After particle inspection and extraction, a total of 721,361 initial particles were used in 2D classification. The best 2D classes were then selected as templates for template picking of particles. Then, a total of 2,134,610 particles were extracted and subjected to 2D classification, yielding a particle set of 1,418,167 particles. After initial reconstruction and heterogeneous refinement with 4 classes, one 3D class accounting for 534,058 particles was generated and refined. The refined particles stack was re-extracted for further heterogeneous refinement with 4 classes, among which two classes (435,496 particles) were combined for non-uniform refinement. Based on this EM map, a mask encompassing one Bre1 dimer and less than half of nucleosome was generated. After C2 symmetry expansion of particles, a total of 870,992 particles were subjected to 3D classification with the above mask. Among the ten 3D classes, five classes had well-resolved Bre1 dimer, and three states of Bre1 binding to nucleosome were identified. The three state maps were further refined to a resolution of 3.47 Å for state 1, 3.25 Å for state 2, and 3.21 Å for state 3. Finally, a sharpening B factor of -5 Å^2^ was applied to maps of state1, 2 and 3. We note that, although Rad6 was fused to the Bre1 C-terminus in an effort to overcome the low affinity of Rad6 for the Bre1 RING domain^12,13^, there was no density corresponding to Rad6 in any of the maps.

PDB structures of Bre1 (PDB: 4R7E) and nucleosome (PDB: 6NOG) were rigid-body fitted to the map using UCSF ChimeraX^25^. The initial Bre1-nucleosome structures were manually rebuilt and subjected to real-space refinement using *Coot*^26^ and Phenix^27,28^. Due to the C2 symmetry of the complex, only one Bre1 dimer was modeled in the final structures. These coordinates were further validated using Comprehensive validation (CryoEM) in Phenix before deposition in the Protein Databank (PDB). Figures were created using PyMOL(Schrödinger), UCSF Chimera, and UCSF ChimeraX.

### Nucleosome ubiquitination assay

For nucleosome ubiquitination reactions, 1 µM NCP, 200 nM E1, 40 µM ubiquitin, 5mM ATP, 5mM MgCl_2_ were mixed with fused E3-E2 (5 µM Bre1-Bre1-Rad6) or separated E3 and E2 enzymes (10 µM Bre1, 5 µM Rad6) in a reaction volume of 20 µL in 20 mM HEPES pH 7.5, 50 mM NaCl, 0.1 mM DTT. Reactions were stopped by addition of 4× SDS loading buffer at indicated time points and analyzed by SDS/PAGE, followed by immunoblotting with Ubiquityl-histone H2B K120 antibody (1:2500 dilution, CST #5546).

### Data availability

The maps are available in the Electron Microscopy Data Bank (EMDB) database under accession No. EMD-41016 (state 1), EMD-41015 (state 2), and EMD-41011 (state 3). The atomic models are available in the Protein Databank (PDB) database with PDB ID 8T3Y (state 1), 8T3W (state 2) and 8T3T (state 3).

